# Generation and characterization of a laforin nanobody inhibitor

**DOI:** 10.1101/2021.01.20.426524

**Authors:** Zoe R. Simmons, Savita Sharma, Jeremiah Wayne, Sheng Li, Craig W. Vander Kooi, Matthew S. Gentry

## Abstract

Mutations in the gene encoding the glycogen phosphatase laforin result in the fatal childhood epilepsy Lafora disease (LD). A cellular hallmark of LD is cytoplasmic, hyper-phosphorylated, glycogen-like aggregates called Lafora bodies (LBs) that form in nearly all tissues and drive disease progression. Additional tools are needed to define the cellular function of laforin, understand the pathological role of laforin in LD, and determine the role of glycogen phosphate in glycogen metabolism. We present the generation and characterization of laforin nanobodies. We identify multiple classes of specific laforin-binding nanobodies and determine their binding epitopes using hydrogen deuterium exchange (HDX) mass spectrometry. Further, one family of nanobodies is identified that serves as an inhibitor of laforin catalytic activity. The laforin nanobodies are an important set of tools that open new avenues to define unresolved questions.

## 1. Introduction

Glycogen is the storage form of glucose and a highly important substrate for cellular metabolism^1^. Nearly all tissues metabolize glycogen and recent work has focused on the importance of glycogen metabolism in the brain^2^. Characterization of the enzymes and mechanisms of glycogen metabolism began over 70 years ago and include: synthesis by glycogen synthase, branching via glycogen branching enzyme, hydrolysis by glycogen phosphorylase, and debranching via glycogen debranching enzyme^3^. Over the last 20 years, a previously unknown protein called laforin has emerged as an important contributor to glycogen metabolism homeostasis.

Multiple labs demonstrated that laforin is a glycogen phosphatase and mutations in the gene encoding laforin cause the formation of aberrant glycogen-like aggregates called Lafora bodies (LBs)^4–6^. LBs are cytoplasmic, water-insoluble aggregates more similar to plant starch than human glycogen^7–10^. The LBs form in cells from nearly all tissues and are the pathological agent driving neurodegeneration and death in Lafora disease (LD) patients^11–14^. LD is autosomal recessive and classified as both a progressive myoclonus epilepsy and glycogen storage disease. The direct relationship between mutated laforin, LB formation, and LB neurotoxicity highlights the importance of glycogen metabolism in the brain. While laforin removes phosphate from glycogen, how phosphate impacts normal glycogen homeostasis or how hyper-phosphorylation is detrimental to glycogen homeostasis and LB formation remains under investigation.

Based on sequence and structural conservation, laforin is classified in the protein tyrosine phosphatase superfamily and the dual specificity phosphatase clade^15^. Laforin is the only known human glycogen phosphatase and is comprised of a dual-specificity phosphatase (DSP) domain and a carbohydrate binding module (CBM). Like all DSPs, laforin contains a protein-tyrosine phosphatase catalytic loop that includes a CX_5_R motif (**C**NAGVG**R**, residues 266–272) at the base of the active site^16^. The architecture and depth of this active site allow laforin to dephosphorylate glycogen^15,17,18^. The laforin CBM belongs to CBM family 20 and allows laforin to bind glycogen as well as other complex carbohydrates *in vitro* and *in vivo*, like LBs, amylopectin, and maltodextrins^8,18,19^. Key structural aspects of laforin that promote glucan-specific phosphatase activity are: closely integrated CBM-DSP domains, an anti-parallel dimerization, and a signature DSP sequence within the active site channel^15^.

In addition to being a glycogen phosphatase, laforin interacts with several proteins involved in glycogen metabolism, including the E3 ubiquitin ligase malin^20^. Approximately 50% of LD patients have mutations in the gene encoding malin and ~50% have mutations in the gene encoding laforin. Malin polyubiquitinates a number of proteins involved in glycogen metabolism, including both laforin and glycogen phosphorylase^20–22^. The consequences of malin-directed ubiquitination are still being elucidated. LD mouse models with a mutation in either gene form hyperphosphorylated LBs and display neurodegeneration^5,23^. However, the relationship between phosphorylation and LD progression remains an unresolved and critical issue^24–26^.

Recombinant antigen-specific, single-domain antibodies, often referred to as nanobodies (Nb), are increasingly used for modulating protein properties for the purpose of elucidating function^27^. While canonical antibodies are comprised of two heavy chains and two light chains, nanobodies are derived from the antigen binding domain of heavy-chain antibodies, produced in camelids and select other organisms. Thus, nanobodies are monomers with a dedicated variable domain, also referred to as VHH, and are ten times smaller than a canonical IgG, i.e. ~15 kDa. Their small size promotes their high propensity for binding unique structural features of proteins, otherwise unreachable by conventional immunoglobulins and capable of manipulating protein conformation linked to function^27–29^. Nanobodies contain three complementary determining regions (CDRs) that are primary determinants of specific antigen binding. Similar affinities between VHHs and conventional antibodies is generally attributed to elongated CDR3 loops in VHH domains, which provide a paratope formed by a single entity^30^.

In this study, we present the characterization of six laforin nanobodies, consisting of five unique CDR3 regions. Two llamas were immunized with recombinant human laforin, 74 potential VHH anti-laforin candidates were screened, and six stably binding anti-laforin nanobodies were characterized. Epitope mapping of the six laforin nanobodies established three general epitopes, one that spans both the CBM and DSP on the opposite side of the DSP active site, one that lies on the CBM, partially covering regions that bind glycogen, and one that covers the DSP active site. Secondary structure epitope mapping established that nanobody Nb72 binds the PTP-loop of the laforin DSP domain. Utilizing three *in vitro* assays, we demonstrate that Nb72 inhibits the phosphatase activity of laforin.

## 2. Methods

### 2.1 VHH library generation

#### 2.1.1 Purification of Laforin

*H. sapiens* (Hs) Laforin residues 1-328 was expressed from pET28b (Novagen) as an N-terminal His6 tagged protein, as previously described^31^. Briefly, laforin was expressed in BL21 (DE3) (Novagen) *E. coli* cells grown in 2xYT media at 37° C until OD_600_ = 0.6, culture flasks were placed on ice for 20 min, induced with 1 mM (final) isopropyl thio-ß-D-galactopyranoside (IPTG), grown for an additional 14h at 20° C, and harvested by centrifugation. Cells were resuspended and lysed in buffer A (20 mM Tris-HCl, 100 mM NaCl, 10% glycerol, 2 mM DTT, pH 7.5), centrifuged, and the proteins were purified using a Profinia immobilized metal affinity chromatography (IMAC) column with Ni^2+^ beads (Bio-Rad) and a Profinia protein purification system (Bio-Rad) using wash (buffer A) and elution buffer (300mM imidazole, 20 mM Tris-HCl, 100 mM NaCl, 10% glycerol, 2 mM DTT, pH 7.5). The desalted elution fraction was further purified using fast protein liquid chromatography (FPLC) with a HiLoad 16/60 Superdex 200 size exclusion column (GE Healthcare). The buffer used for laforin purification for the small-scale pulldowns described in methods 2.3.2 was (50 mM HEPES, 100 mM NaCl, 10% glycerol, 2 mM DTT, pH 7.5. For all other experiments, laforin was purified in 20 mM Tris-HCl, 100 mM NaCl, 10% glycerol, 2 mM DTT, pH 7.5.

#### 2.1.2 Immunizations

Inoculation, construction of the VHH libraries, panning, and cloning was performed by the VIB Nanobody Core (Vrije Universiteit Brussel, Brussels). Two llamas were subcutaneously injected on days 0, 7, 14, 21, 28 and 35 with 250 μg/animal of recombinant laforin emulsified with Gerbu adjuvant P. On day 40, anticoagulated blood was collected for lymphocyte preparation.

#### 2.1.3 Construction of the VHH libraries

A VHH library was constructed from each llama to screen for the presence of laforin specific nanobodies. First strand cDNA synthesis was achieved using an oligo(dT) primer and total RNA from peripheral blood lymphocytes as a template. The VHH encoding cDNA sequences were amplified by PCR, digested with PstI and NotI, and cloned into the PstI and NotI sites of the phagemid vector pMECS.

#### 2.1.4 Isolation of laforin specific nanobodies

Each library was panned individually for three rounds on solid-phase coated antigen (200 μg/ml in 100 mM NaHCO_3_, pH 9.3). After each round of panning, the enrichment for laforin-specific phages was assessed by comparing the number of phagemid particles eluted from antigen-coated wells with the number of phagemid particles eluted from negative control (uncoated blocked) wells. In total, 950 colonies (475 colonies for each library: 95 from panning round one, 285 from round two, and 95 from round three) were randomly selected and analyzed by ELISA for the presence of laforin specific nanobodies in their periplasmic extracts. The antigen used for panning and ELISA screening was the same as the one used for immunization, using uncoated blocked wells as negative control. Out of these 950 colonies, 138 colonies scored positive in this assay. Based on sequence data of the positive colonies, 74 different full-length nanobody candidates were distinguished based on complementary determining regions.

#### 2.1.5 Sequence verification, alignment, and evolutionary analysis

Nanobody sequences were verified using the primer: 5’ TTA TGC TTC CGG CTC GTA TG 3’ and the six direct laforin binding nanobody plasmids will be deposited to https://www.addgene.org/. All sequence alignments were performed using Clustal-Omega available on the European Bioinformatics Institute’s server http://www.ebi.ac.uk/. The evolutionary tree was calculated from the alignment using neighbor-joining clustering implemented in the Clustal-Omega package^32^. The guide tree was displayed with the program iTOL (http://itol.embl.de/)^33^.

### 2.2 Screening for VHH expression

74 potential VHH candidates were transformed into BL21 (DE3) *E. coli* cells (MilliporeSigma) and expression levels were analyzed by Western analysis. Nanobodies were expressed from pMECS as a C-terminal HA and His6 fusion in 10 mL cultures grown in 2xYT until OD_600_ = 0.6. 100 μL of each culture was aliquoted, centrifuged, air dried, and stored at −20° C while protein expression was induced in the remaining cultures using 1 mM IPTG (final concentration). Induced cultures grew for 3.5h, 80 μL of the induced cultures were pelleted, air dried and stored at −20° C. The 100 μL uninduced and 80 μL induced culture pellets were lysed with 75 μL of 50 mM TRIS, 8M urea by vortexing. Total proteins from the bacteria culture lysate were resolved by stain-free gel SDS-PAGE (Bio-Rad). Nanobody expression was visualized by Western analysis using an anti-6xHis antibody (1:1000) (NeuroMab #75-169) and goat anti-mouse secondary (1:3000) (Invitrogen **#**62-6520).

### 2.3 Small scale pulldowns

#### 2.3.1 Small scale nanobody purification

Nanobodies screened for expression were purified using the remaining 9.82 mL of induced culture, which was pelleted and frozen at −20° C. Nanobodies were purified from the *E. coli* pellets with Ni-NTA resin. *E. coli* cell lysates were centrifuged to separate soluble protein which was incubated with Ni-NTA agarose beads for 1 h at 4° C in 1 mL bind buffer (20 mM Tris-HCL, 100 mM NaCl, 15 mM Imidazole, pH 8.0). The samples were washed three times in bind buffer and eluted in 100 μL of 300 mM imidazole, 100 mM NaCl, pH 8.0 and used for small scale pulldowns.

#### 2.3.2 Antigen affinity pulldown

Primary screening for nanobody binding was achieved by a small-scale pulldown method in which Affi-gel 10 affinity resin (Bio-Rad) was bound to laforin and subsequently to each purified VHH clone^34^. Expression and purification of laforin was performed as described above in 50 mM HEPES, 100 mM NaCl, 10% glycerol, 2 mM DTT, pH 7.5. 50 μL Affi-gel (100 μL slurry) per pulldown was washed and resuspended in cold PBS per manufacturer’s directions. The washed Affi-gel was saturated with laforin in 50 mM HEPES, 100 mM NaCl, 10% glycerol, 2 mM DTT, (final volume 500 μL) and incubated with rocking for 1 h at 4° C. The laforin bound Affi-gel resin was spun down at 300 rpm for 2 min and washed with cold PBS two times. Affi-gel alone was used as a control. Then, the laforin bound or unbound Affi-gel was blocked with cold 1X TBS for 1 h at 4° C. The samples were spun down, supernatant was removed, and 75 μL (~4-12 μg) of small batch Ni^2+^ purified nanobodies were added. Pulldowns were washed three times with 1X PBS and the supernatant was removed. Bound proteins were denatured in Laemmli’s buffer and ~12 μL of each sample was loaded per well of a stain free SDS-PAGE gel and imaged.

### 2.4 Nanobody purification

Larger concentrations of purified nanobodies were achieved by scaling up methods described in 2.2 and 2.3.1. Nanobodies were expressed in BL21 (DE3) *E. coli* cells grown in 2xYT at 37° C to OD_600_ = 0.9. 15 mL of pre-culture was used per L culture. Cultures were induced with 1 mM IPTG (final) and incubated for ~16-18 h at 25° C after which, pellets were harvested by centrifugation. Nanobodies were purified from the *E. coli* pellets with Ni-NTA resin. *E. coli* cell lysates were centrifuged to separate soluble protein which was incubated with Ni-NTA agarose beads for 1 h at 4° C in 30 mL bind buffer (20 mM Tris-HCL, 100 mM NaCl, 15 mM Imidazole, pH 8.0). The samples were washed three times in bind buffer and eluted in 300 mM imidazole, 100 mM NaCl, pH 8.0. Eluted samples were separated from the Ni-NTA beads by filtering through a 25 mM syringe filter and then buffer exchanged with 20 mM Tris-HCL pH 7.5, 100 mM NaCl using FPLC with a HiLoad 16/60 Superdex 75 column (GE Healthcare).

### 2.5 Size exclusion analysis of VHH-laforin complexes

Laforin and VHH were combined in molar ratio (~1:1.1) and resolved by size exclusion chromatography (SEC) using a HiLoad 10/300 Superdex 200 size exclusion column and 100 mM NaCl, 20 mM TRIS, 10% glycerol, 0.35 mM β-mercaptoethanol (BME), pH 7.5. WT laforin alone was utilized as a control. Fractions were collected in 500 μL volumes and visualized by loading 20 μL of each fraction onto a stain-free gel SDS-PAGE (Bio-Rad) and imaged.

### 2.6 Hydrogen Deuterium Exchange Mass Spectrometry (HDX)

HDX experiments were performed as previously described^18,35^. The optimal peptide coverage map of laforin was obtained using a quench solution containing 0.08 M GuHCl, 0.1 M Glycine, 16.6% Glycerol, pH 2.4. To initiate HDX experiments, 3 μL of stock solution (laforin or laforin-nanobody complex at 1 mg/ml) was mixed with 9 μL of D_2_O buffer (8.3 mM Tris, 50 mM NaCl, pD_READ_ 7.2) and incubated at 0° C for 10, 100, 1000 and 10,000 sec. The exchange reaction was quenched by adding 18 μL of the above quench solution and the quenched samples were flash frozen with dry ice. All frozen samples, including un-deuterated and equilibrium-deuterated control samples, were passed over an immobilized pepsin column (16 μL) at a flow rate of 25 μL/min and digested peptides were collected on a C18 trap column (Optimize Tech, Opti-trap, 0.2×2 mm) for desalting. The peptide separation was performed on a C18 reverse phase column (Agilent, Poroshell 120, 0.3×35 mm, 2.7 μL) with a linear gradient of 8-48% B over 30 min (A: 0.05% TFA in H_2_O; B: 80% acetonitrile, 0.01% TFA, and 20% (H2O)). Mass spectrometry (MS) analysis was performed on the Orbitrap Elite mass spectrometer (Thermo Fisher Sci), which was adjusted for HDX experiments^36^. The resolution of the instrument was set at 120,000 at m/z 400. Proteome Discoverer software (v1.3, Thermo Scientific) was used to identify the sequence of the digested peptide ions from their MS/MS data. HDXaminer (Sierra Analytics, Modesto, CA) was utilized to confirm the peptide identification and calculate the centroids of isotopic envelopes of all the peptides. The level of deuterium incorporation of each peptide was calculated by applying back-exchange correction^37^. The ribbon maps were generated from deuteration level of overlapping peptides to improve the resolution of the HDX data.

### 2.7 Phosphatase assays

#### 2.7.1 *para-*Nitrophenyl Phosphate (pNPP) assay

Generic phosphatase activity assays were performed using *para*-Nitrophenyl Phosphate (pNPP) as previously described^4,38^. Assays were performed using 96 well plates in 50 μL reactions containing 1X phosphatase buffer (0.1 M sodium acetate, 0.05 M Bis-Tris, 0.05 M Tris-HCl, pH 5.0), 2 mM dithiothreitol (DTT), and 50 mM pNPP. To maintain reactions in the linear phase, 250, 500, or 1000 ng of laforin pre-bound to an equal molar quantity of nanobody were added to the reaction in triplicates and incubated in a 37° C water bath for 10 minutes. The reaction was terminated by the addition of 200 μL of 0.25 M NaOH and absorbance was measured at 410 nm.

#### 2.7.2 Glucan dephosphorylation assays

Glucan phosphatase activity assays were performed as previously described^4,38^. Assays were performed using 96 well plates in 100 μL reactions containing 1X phosphatase buffer (100 mM sodium acetate, 50 mM bis-Tris, 50 mM Tris-HCl, pH 7.0) and 2 mM DTT, and 10mg/mL rabbit skeletal muscle glycogen or 1 mg/mL potato amylopectin (MilliporeSigma). To maintain enzymatic activity in the linear phase, every 10 seconds, 250, 500, or 1000 ng of laforin pre-bound to an equal molar quantity of nanobody were added to the reaction mixture in quadruplicates and incubated for 30 min at RT. The reaction was stopped by the addition of 25 μL of malachite gold reagent mix added every 10 seconds. Once each well was given the malachite gold reagent mix, stabilizer was added to each well in intervals of 10 seconds. Absorbance was measured at 635 nm.

#### 2.7.3 Statistical Analysis

Two-way ANOVA was performed using GraphPad Prism version 9.0.0 for Mac, GraphPad Software, San Diego, California USA, www.graphpad.com

## 3. Results

### 3.1 Anti-Laforin nanobody primary screen

Two llamas were immunized with purified recombinant human laforin. Lymphocytes were isolated and the VHH coding sequences were cloned into a phagemid vector. After three rounds of panning and ELISA screens using recombinant laforin protein, a total of 74 clones scored positive and were sequenced, revealing 37 different CDR3 groups (**Figure S1**). A sequence similarity tree reveals the presence of two nanobody groups in which 35 clones are found in one and 39 are found in the other (**Figure 1**). All 74 clones were transformed into BL21 *E. coli* cells and VHH expression was determined by western analysis using an anti-HIS antibody. Approximately 50% of the clones were eliminated due to lack of expression in BL21 cells (data not shown). The expressing clones were each purified on a small scale and screened for their capacity to bind laforin by a pulldown method using affinity resin.

**Figure 1.**
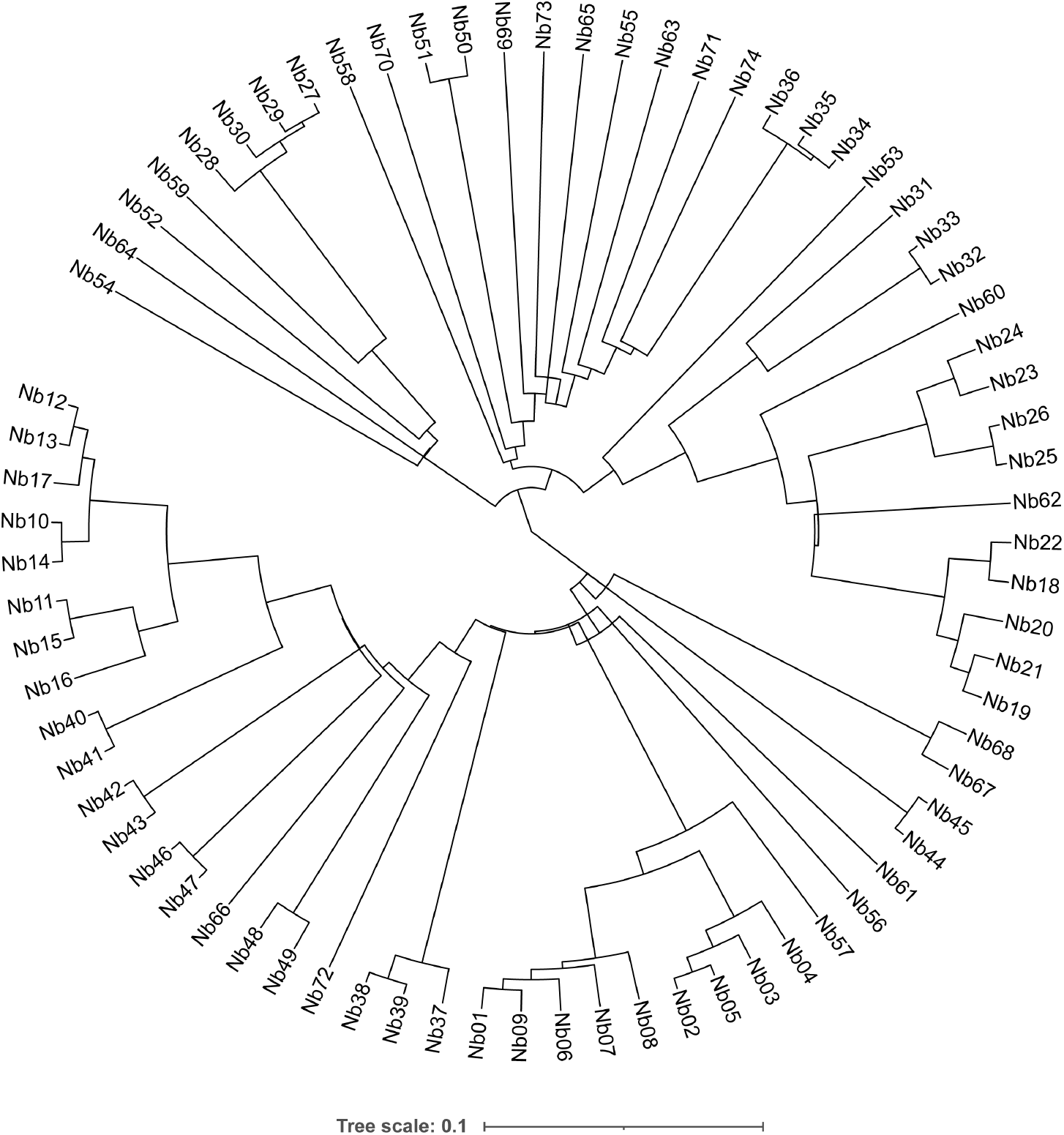
Anti-laforin nanobody guide tree. A binary tree wasgenerated from a sequence alignment and subsequent tree generation of the 74 generated laforin nanobody clones using Clustal-Omega.

To identify nanobodies that form a stable complex with laforin, the ability of purified nanobodies to be precipitated by laforin affinity resin was assessed. Laforin coupled resin was incubated individually with each of the purified VHH clones, the resin was washed, and bound proteins were eluted and analyzed by SDS-PAGE. This primary screen identified six clones as having direct antigen binding (**Figure 2A**).

**Figure 2.**
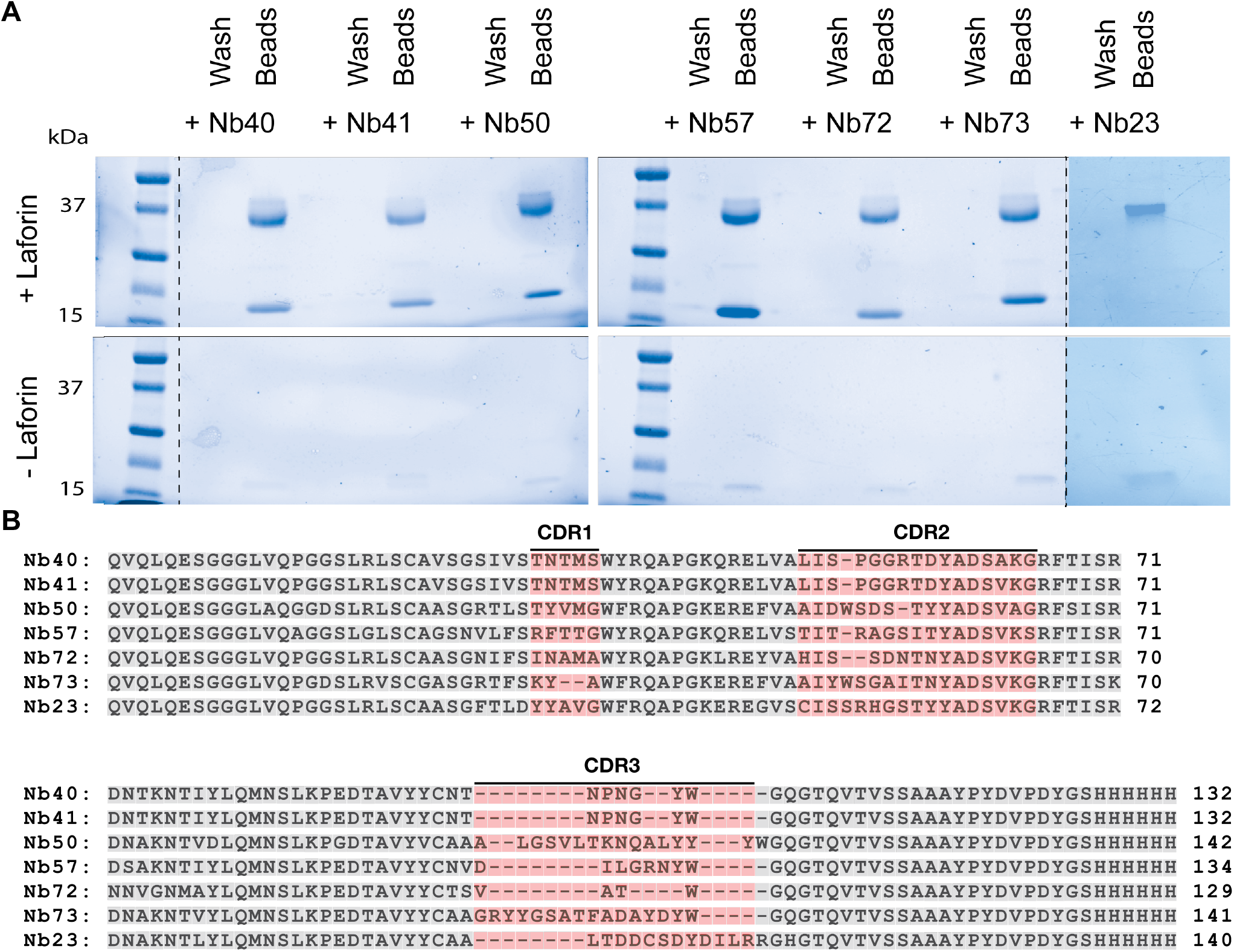
Direct antigen binding nanobodies and comparison of their sequences. **(A)** Primary screening for nanobody binding was achieved by incubating laforin with affinity resin, washing the resin, blocking the resin with TBS, incubating the laforin bound affinity resin with a nanobody, and washing the resin before the laforin-nanobody complex was eluted. Free (wash) and bead bound laforin and/or nanobody was loaded onto an SDS-PAGE stain-free gel and imaged (top). As a control, affinity resin without bound laforin was blocked and then incubated with nanobody (bottom). **(B)** Sequence alignment of nanobodies in A. Sequence homology is 55.9% identical. Complementary determining regions are highlighted in red and framework regions are highlighted in gray (Kabat).

The six identified anti-laforin nanobodies are: Nb40, Nb41, Nb50, Nb57, Nb72, and Nb73. Based on phylogenetic analysis and sequence alignment (**Figure 1 and Figure 2B**), the six identified nanobodies that bind laforin exhibit sequence diversity and contain five different CDR3 regions. Complete DNA sequences of the six laforin nanobodies are provided in **Table S1**.

### 3.2 Size-exclusion analysis of anti-laforin nanobodies complexed to laforin

Size-exclusion chromatography (SEC) was utilized to detect and characterize the laforin-nanobody complexes. Each nanobody was incubated with laforin in ~10% molar excess, samples were subjected to SEC, fractions were collected, and analyzed by SDS-PAGE (**Figure 3**). The laforin-nanobody complexes eluted between 12.11-14.51 mL, laforin alone eluted at 15.40 mL, and the nanobodies alone eluted between 18.08-19.53 mL. All six laforin-nanobody complexes eluted prior to laforin or the nanobody alone, indicating a higher molecular weight and supporting the results of the primary screen. Similarly, SDS-PAGE analyses of the fractions demonstrate co-elution of the laforin-nanobody complexes. Nb40 and Nb73 complexed with laforin eluted earlier than others (**Figure 3**). These supershifted complexes may indicate an increase in nanobody to laforin stoichiometry. Alternatively, Nb40 and Nb73 may interfere with the laforin CBM binding to the carbohydrate-based resin. Laforin CBM mutations have been shown to interrupt this interaction and change the elution time^39^.

**Figure 3.**
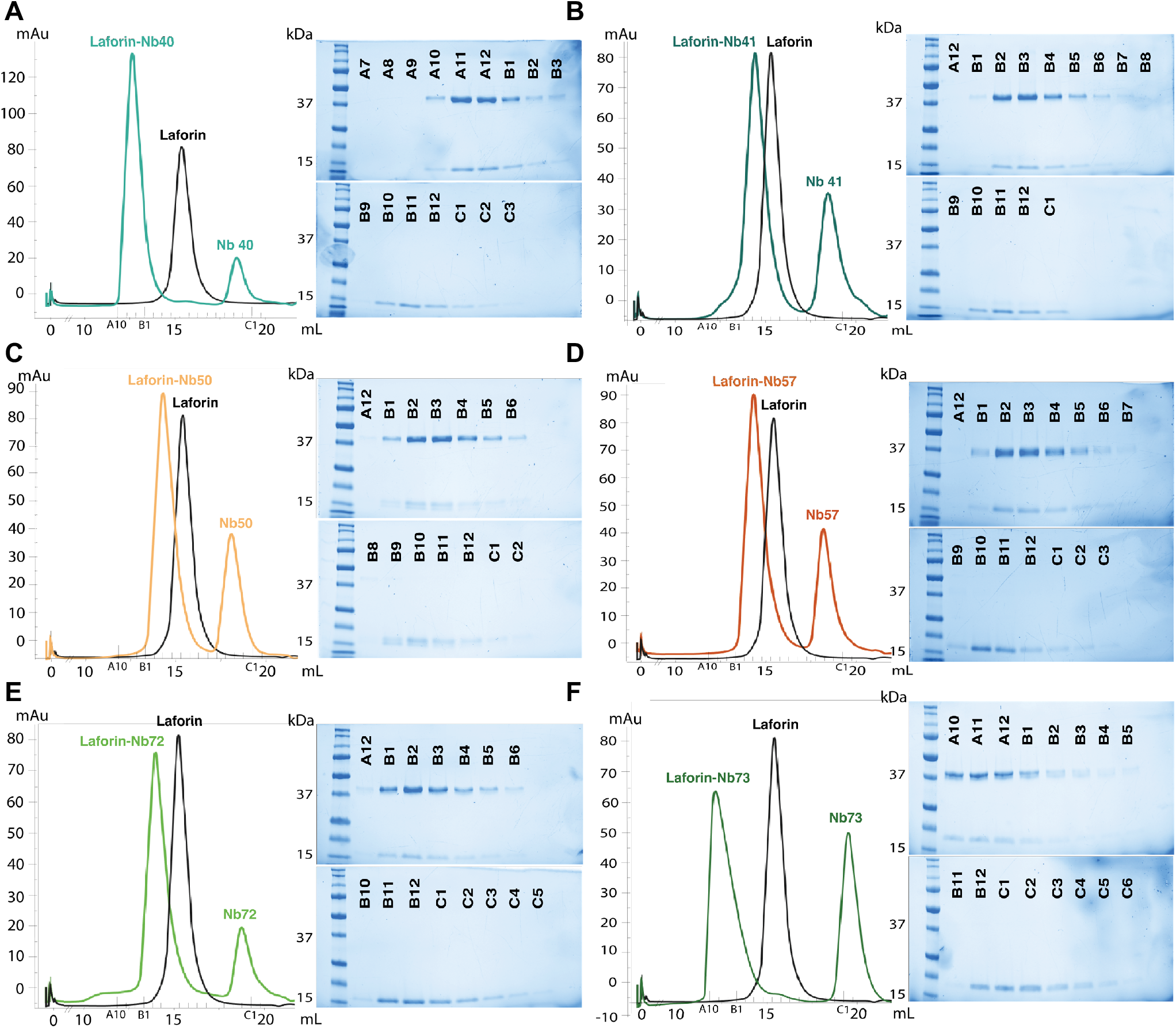
Size-exclusion (SEC) analysis of anti-laforin nanobodies complexed to laforin. For each SEC analysis of the laforin-nanobody complex, the respective fractions are visualized by an SDS-PAGE stain-free gel shown to the right. **A-F)** SEC analysis of laforin alone is shown in black (15.4mL). **A)** Nb40 (18.45mL) complexed to laforin in light teal (12.65mL). **B)** Nb41 (18.58mL) complexed to laforin in dark teal (14.51mL). **C)** Nb50 (18.08mL) complexed to laforin in light orange (14.28mL). **D)** Nb57 (18.19mL) complexed to laforin in dark orange (14.3mL). **E)** Nb72 (18.89mL) complexed to laforin in light green (14.12mL). **F)** Nb73 (19.53mL) complexed to laforin in dark green (12.11mL).

### 3.3 Nanobody epitope mapping via Hydrogen–Deuterium Exchange Mass Spectrometry (HDX)

Epitope mapping can be accomplished using hydrogen-deuterium exchange (HDX) mass spectrometry. HDX quantifies protein surfaces that are solvent accessible. When a nanobody binds a protein, the surface of the protein becomes less solvent accessible. This decrease in solvent accessibility can be quantified by HDX.

Each laforin-nanobody complex and laforin alone were incubated in deuterated buffer for 10, 100, 1,000, and 10,000 seconds (**Figure 4 and S2**). A decrease in laforin deuteration (≥15% decrease in deuteration compared to WT) emerged with Nb40 and Nb41 (**Figure 4A and S2A**), Nb50 and Nb57 (**Figure 4B and S2B**), and Nb72 and Nb73 (**Figure 4C and S2C**). Since Nb40 and Nb41 differ by only one amino acid, their shared epitope is expected. Their epitope partially maps across the CBM (residues 60-66, 120-124, and 126-129) with the majority spanning across the recognition domain of the DSP (residues 139-145, 148-161, 163-169, and 171-186). The epitope of Nb41 extends further on the DSP (residues 193-197). Sequence homology between Nb40 and Nb41 is 99.2% (**Figure S3**). Nb50 and Nb57 share a similar epitope that encompasses most of the CBM (residues 30-33, 35-41, 43-52, 60-66, 120-124) and partially across the DSP (residues 287-291). Nb57 extends further on the CBM (residues 43-56, 58-66, and 126-129). Sequence homology between Nb50 and Nb57 is 66.2% (**Figure S3**). Nb50 and Nb57 share a partial CBM epitope with Nb40 and Nb41. The DSP epitope of Nb40 and Nb41 partially overlaps with Nb72 and Nb73. The epitopes of Nb72 and Nb73 are mapped solely on the DSP and overlap with the recognition domain (residues 139-145, 148-155, 193-197, 236-242). Importantly, Nb72 includes the PTP-loop (residues 267-275). Intriguingly, the sequence homology between Nb72 and Nb73 is 70.6% and no sequence homology exists between their CDR3 regions (**Figure S3**).

**Figure 4.**
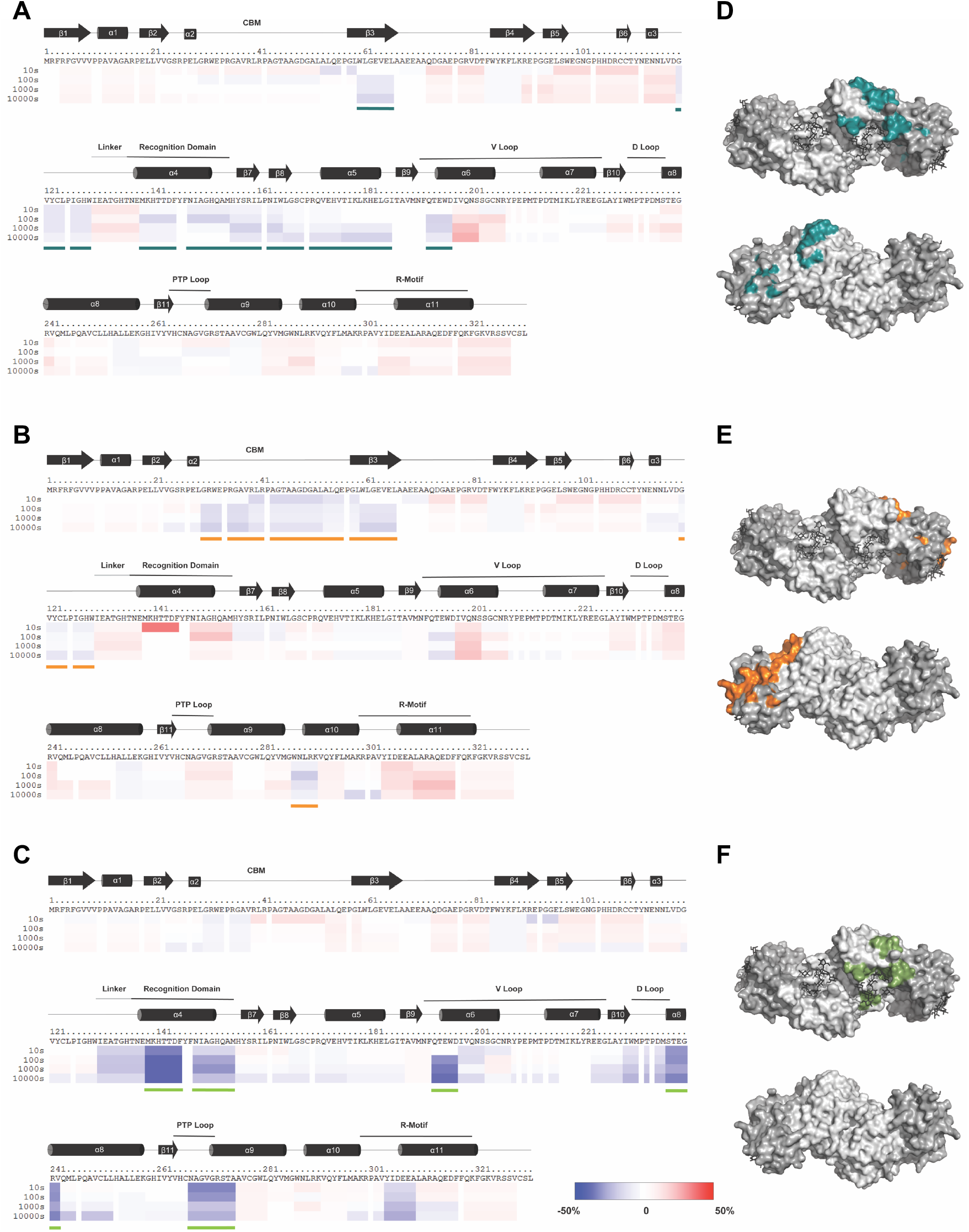
HDX analysis of anti-laforin nanobody binding. Fractions of each nanobody bound to laforin (Figure 3) were analyzed by hydrogen deuterium exchange mass-spectrometry (HDX) and the deuteration level was compared to laforin alone. The percent difference in deuteration between the nanobody complex versus laforin alone is represented with a negative percent change in blue to a positive percent change in red. Each colored bar below the primary sequence represents percent change in deuteration at one of four time points. Secondary structure elements are depicted above the primary sequence with the CBM and DSP motifs labeled. Regions where the nanobody caused a ≥15% change in deuteration are highlighted with a colored line. **A)** The Nb41 epitope on laforin spans β3, β7 and β8 sheets, α4, α5, and partially α6 helixes. Highlighted with a dark teal line. **B)** The Nb57 epitope on laforin includes the β3 sheet and partially covers the α10 helix. Highlighted with a dark orange line. **C)** The Nb72 epitope on laforin covers the α4 helix, and partially spans the α6, α8, and α9 helixes. Highlighted with a light green line. **D,E,F)** The identified epitope for each nanobody was modeled onto the surface map of the laforin structure (PDB: 4RKK). Nb41 (dark teal), Nb57 (dark orange), and Nb72 (light green) respectively mapped on laforin. The CBM and DSP of laforin are colored dark gray and light gray respectively.

These results indicate that three general binding regions exist among the six laforin nanobodies. Nb40 and Nb41 span both the CBM and DSP on the opposite side of the DSP active site (**Figure 4D and S2D**). Nb50 and Nb57 span the CBM, partially covering regions that bind glycogen (**Figure 4E and S2E**). Nb72 and Nb73 bind a region near the DSP active site and Nb72, but not Nb73, overlaps with the PTP-loop (**Figure 4F and S2F**).

### 3.4 Nb72 inhibits the general phosphatase activity of laforin

Given the binding site of Nb72, we predicted that this nanobody would decrease or inhibit the phosphatase activity of laforin. To assess laforin’s general phosphatase activity, we utilized the phosphatase substrate *para*-nitrophenyl phosphate (pNPP). Protein tyrosine phosphatases, including laforin, can convert pNPP to *para*-nitrophenyl (pNP) that results in a colorimetric change that can be quantified at OD_410_. Three concentrations of recombinant laforin were incubated with a respective and ~1:1 molar ratio of nanobody Nb72 or Nb23, a nanobody that did not co-precipitate with laforin (**Figure 2**). As controls, recombinant laforin was also pre-incubated in buffer lacking a nanobody and nanobodies were tested in the absence of laforin. The pNPP substrate was added to each combination using previously defined optimal conditions, reactions were quenched with sodium hydroxide, and dephosphorylation levels were quantified at OD_410_. Laforin incubated with Nb23 yielded similar levels of pNPP dephosphorylation as laforin alone, indicating that Nb23 did not inhibit laforin phosphatase activity (**Figure 5A**). Conversely, laforin incubated with Nb72 displayed a dramatic 85% reduction in pNPP phosphatase activity (**Figure 5A**).

**Fig 5.**
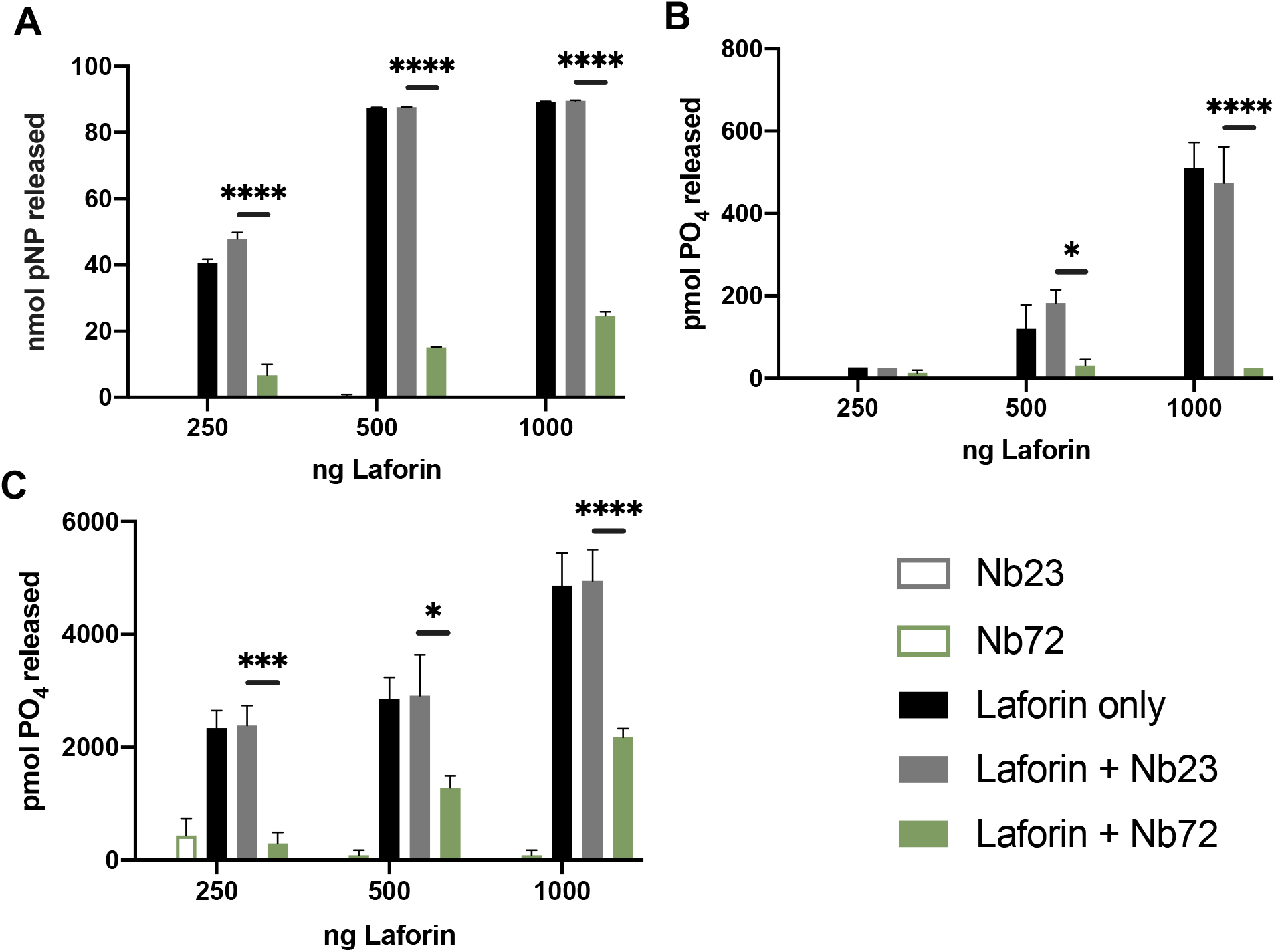
Laforin activity in the presence of Nb72. **A)** The specific activity of laforin against pNPP in the presence and absence of nanobodies. **B)** The specific activity of laforin against rabbit skeletal muscle glycogen in the presence and absence of nanobodies. **C)** The specific activity of laforin against potato amylopectin in the presence and absence of nanobodies. Each bar is the mean ±SEM of 4 replicates. * p < 0.0301, *** p = 0.0007, **** p < 0.0001.

### 3.5 Nb72 inhibits the glycogen phosphatase activity of laforin

The small molecule pNPP is a readout for general phosphatase activity. However, pNPP can integrate into the DSP active site independent of the laforin CBM to be converted into pNP. In cells, laforin binds glycogen via its CBM and engages the phospho-glucose substrate with its DSP active site to dephosphorylate the substrate. Therefore, we sought to determine if Nb72 binding to laforin inhibits its specific glucan phosphatase activity.

Glycogen dephosphorylation can be assessed by quantifying phosphate release using a malachite gold colorimetric assay. Three concentrations of recombinant laforin were incubated with nanobody Nb72 or the negative control nanobody Nb23 in a respective and ~1:1 molar ratio. As a control, recombinant laforin was also pre-incubated in buffer lacking a nanobody. Then, rabbit skeletal muscle glycogen was added as the substrate using previously defined optimal conditions. Reactions were incubated at 20° C for 30 min, and then quenched with the phosphatase inhibitor N-ethylmaleimide. Malachite gold solutions were added, and dephosphorylation levels were quantified at OD_635_. Laforin incubated with Nb72 yielded a dramatic 95% reduction in laforin’s glycogen phosphatase activity while Nb23 did not reduce the activity (**Figure 5B**). No other nanobody showed a significant inhibitory effect on laforin’s phosphatase activity towards amylopectin (**Figure S4**).

Laforin’s endogenous substrate is glycogen^4,6,40^. However, laforin has a higher binding affinity for LBs^19,41^. While glycogen is water-soluble, the glucose chains of LBs are longer and predicted to form helical structures making LBs more water-insoluble^7–9^. Plant amylopectin is also a water-insoluble glucan that has covalently attached phosphate and has been utilized as a LB proxy^4,8,40,42^. Therefore, we performed a similar assay as above utilizing malachite gold, laforin and the nanobodies with amylopectin as the substrate. Overall, laforin released ~10-fold more phosphate from amylopectin than glycogen, which is as expected given the higher level of phosphate in amylopectin^43,44^. Nb72 was observed to significantly inhibit laforin phosphatase activity by 55% against amylopectin (**Figure 5C**). These data further strengthen a growing body of results suggesting that laforin has a higher degree of binding and/or activity towards insoluble substrates.

## 4. Discussion

The glycogen phosphatase laforin is critical for brain glycogen metabolism. Mutations in the laforin gene promote the formation of cytoplasmic, insoluble glycogen-like aggregates that are the pathological agent of LD^6,45,46^. The mechanism by which laforin influences the solubility of glycogen is still unknown and progress in this discovery could be accelerated with additional tools. The current study showcases the generation of six laforin nanobodies, mapping of the nanobody epitopes by HDX, and three *in vitro* assays demonstrating that nanobody Nb72 inhibits laforin’s phosphatase activity.

HDX nanobody epitope mapping revealed that one nanobody, Nb72, decreased deuterium exchange within the laforin PTP-loop, implying that Nb72 could inhibit the laforin phosphatase activity. Three *in vitro* phosphatase activity assays established that binding Nb72 to laforin markedly reduces the phosphatase activity of laforin. The first assay employed a non-specific substrate*, para*-nitrophenyl phosphate (pNPP), to assess the inhibitory activity of Nb72. The pNPP assay quantifies laforin phosphatase activity independent of glycogen binding. Nb72 decreased laforin phosphatase activity by 85%. Using more biologically relevant substrates, we discovered that Nb72 diminishes laforin phosphatase activity against skeletal muscle glycogen by 95%, and amylopectin by 55%. Thus, Nb72 is more effective at inhibiting laforin’s phosphatase activity directed towards glycogen. These data are consistent with published reports demonstrating that laforin preferentially binds more water-insoluble substrates like amylopectin and LBs^19,47^. The reason for this preference has been proposed to be that the CBM domain has an enhanced contribution to phosphatase activity as the carbohydrate increases in complexity and/or insolubility^42^. Laforin is dramatically stabilized in the presence of a longer chain length oligosaccharide compared to a shorter chain length oligosaccharide. Further, the presence of the longer chain length oligosaccharide promotes cooperativity in binding between the dimers of laforin that is not observed with a shorter chain length sugar substrate^18^. These data are further supported by our findings that more phosphate is released from amylopectin than glycogen in the presence of laforin. The ~10-fold difference in phosphate release can be accounted for by the ~10-fold difference in total phosphate contained in amylopectin versus glycogen. Glycogen contains 1 phosphate per 1,000-10,000 glucose residues and amylopectin contains 1 phosphate per ~300 glucose residues^5,42,48–51^.

Nanobodies are a rapidly growing technology that are being utilized in novel ways to refine a wide variety of traditional techniques such as crystallization chaperoning, affinity purification, immunoprecipitation, super-resolution microscopy, confocal microscopy, flow cytometry, cell delivery, radiolabeling, and modulating protein function and interactions in cells^52,53^. An immediate opportunity to utilize the laforin nanobodies is with respect to modulating laforin’s phosphatase activity, glycogen binding, and interactions. Nb72 clearly inhibits the glycogen phosphatase activity of laforin *in vitro*. **Figure S4** points to the possibility that Nb50 and Nb57, CBM-binding nanobodies, may interfere with laforin’s catalytic activity (though the effect is not statistically significant at the given concentration), and may modulate laforin’s carbohydrate binding in cell culture. These nanobodies could be expressed in cells to probe the relationship between glycogen phosphate with glycogen metabolism, central carbon metabolism, and metabolic signaling events without imposing structural mutations to laforin.

Another opportunity to use the laforin nanobodies is protein crystallography. The only full-length laforin structure to date is the catalytically inactive human Laforin-C266S^18^. While this structure has provided key insights, as described above, a structure of the wildtype and patient mutations is needed to fully define the catalytic cycle and resolve why laforin mutations cause LD. This work has been hampered due to the molecular dynamics of wildtype and mutant enzymes. Nanobodies can be employed to limit protein dynamics and there have been impressive successes in this area^52^. Crystallizing laforin with the nanobodies could capture conformations that are important for its biological function.

Multiple groups have demonstrated that laforin and malin physically interact and form a complex^54^. Additionally, the *in vivo* function of laforin likely includes a sophisticated scaffolding and/or signaling role because multiple studies have found that laforin interacts with several glycogen metabolism related proteins^4,55,56^. In fact, many laforin protein interactions are both involved in glycogen metabolism and are putative substrates of malin, including the protein phosphatase 1 regulatory subunits GL, R5 and R6/PTG; glycogen synthase, the pyruvate kinase isoforms PKM1 and PKM2; and AMP-activated protein kinase β subunits^45,54,57,58^. Due to these interactions, the laforin-malin complex has been suggested to participate in protein degradation, oxidative stress, and the unfolded protein response^55^. Therefore, LD pathology could arise from mutations that affect the specific activity of laforin and/or malin, and also from mutations that impair the interaction of both proteins to form a complex. Select nanobodies could be utilized to further elucidate these intercalated signaling events. Further defining which nanobodies block laforin’s protein interactions and expressing those nanobodies in cell culture may illuminate the mechanism of laforin’s role in normal glycogen metabolism and LD suppression. Cellular expression of the laforin nanobodies has the potential to block laforin-protein interactions and greatly increase the field’s understanding of the mechanics and purpose of the laforin-malin complex, both in disease and normal cellular metabolism.

## Acknowledgments

This work was supported by National Institutes of Health grants NS097197 and NS116824 (to M.S.G.) and TR001997 (to Z.R.S). The content is solely the responsibility of the authors and does not necessarily represent the official views of NIH. We wish to thank Dr. Ramon Sun and the Sun lab for useful advice as well as VIB Nanobody Core (Vrije Universiteit Brussel, Brussels) for work in the initial VHH library generation. This paper is subject to the NIH Public Access Policy. This study was carried out in accordance with the Uniform Requirements for Manuscripts Submitted to Biomedical Journals.

## Corresponding Author

Dr. Matthew S. Gentry, University of Kentucky, College of Medicine, Department of Molecular and Cellular Biochemistry, Center for Structural Biology, 741 S. Limestone, RM B177, Lexington, KY 40536-0509, Phone: +001 (859) 323-8482.

## Author contribution

Z.R.S performed the experiments, analyzed the experimental data, generated the figures, and wrote the manuscript. S.S. assisted in experimental design and in interpreting the experimental data. J.W. performed nanobody purifications. S.L. performed the HDX experiments. C.V.K. assisted in the conception of the project, data interpretation, and manuscript preparation. M.S.G. conceived the project, aiding in experimental planning, analysis of the experimental data, and preparation/revision of the manuscript.

## Abbreviations

LB: Lafora body
LD: Lafora disease
DSP: Dual-specificity phosphatase
CBM: Carbohydrate binding module
Nb: Nanobody
VHH: Single-domain heavy chain antibodies
CDR: Complementary determining regions
PTP: Protein tyrosine phosphatase
Laforin: Recombinant human N-terminal His6 Laforin 329X
IPTG: Isopropyl thio-ß-D-galactopyranoside
IMAC: Immobilized metal affinity chromatography
FPLC: Fast protein liquid chromatography
HEPES: N-2-hydroxyethylpiperazine-N-ethanesulfonic acid
DTT: Dithiothreitol
PCR: Polymerase chain reaction
ELISA: Enzyme-linked immunosorbent assay
SDS-PAGE: Sodium dodecyl sulfate-polyacrylamide gel electrophoresis
Ni-NTA: Nickel-nitrilotriacetic acid (nickel-charged affinity resin)
PBS: Phosphate-buffered saline
SEC: Size-exclusion chromatography
BME: β-mercaptoethanol
HDX: Hydrogen-deuterium exchange mass spectrometry
TFA: Trifluoroacetic acid
MS: Mass spectrometry
pNPP: *para*-Nitrophenyl Phosphate
pNP: *para*-Nitrophenyl
DNA: Deoxyribonucleic acid
AMP: Adenosine monophosphate

## Supplemental Figures

**Figure S1.**
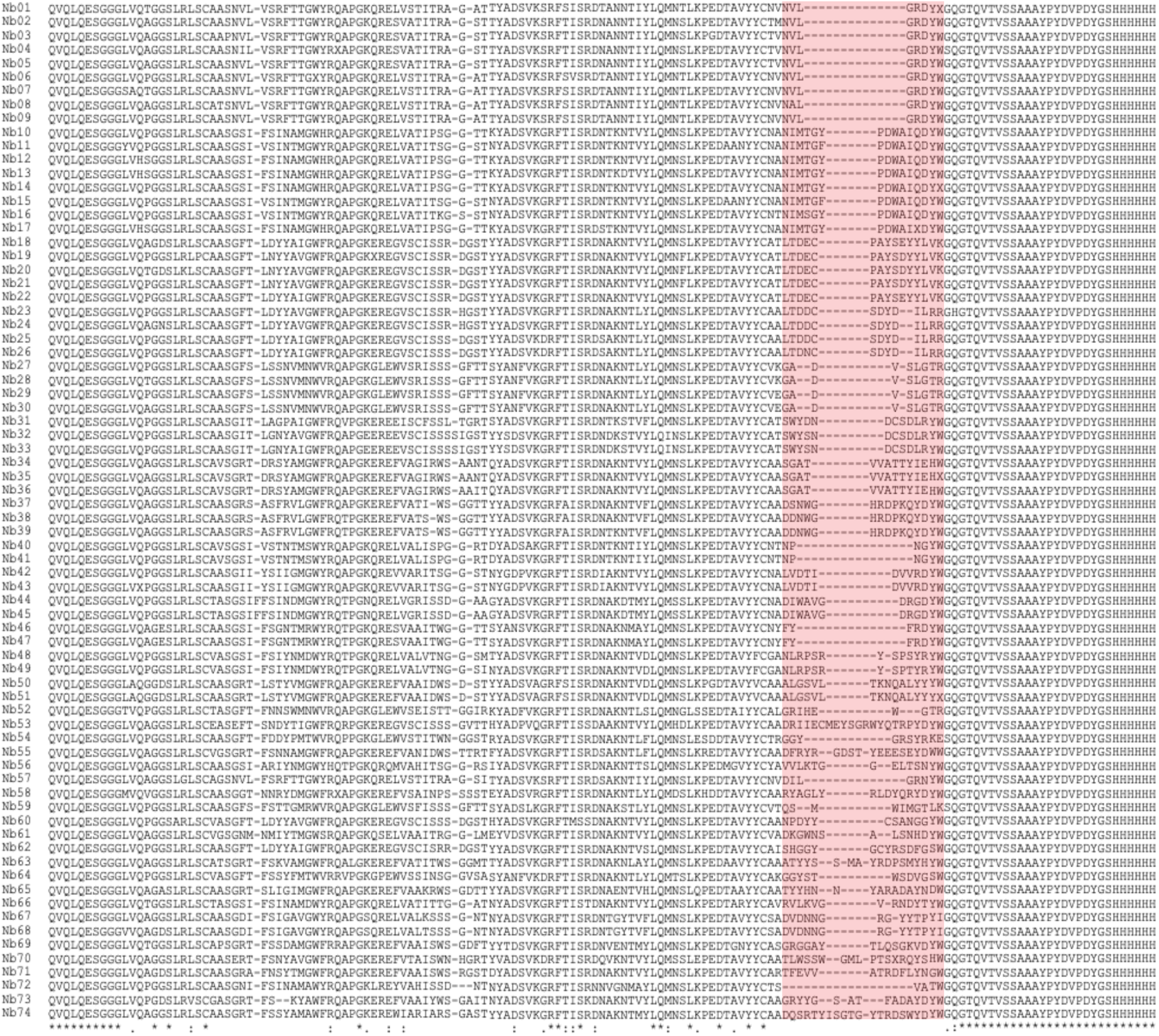
Anti-laforin nanobody sequence alignment. 37 CDR3 groups are represented among the 74 generated laforin nanobodies (highlighted in red). Sequence homology of the 74 nanobodies is 33.1% (Clustal-Omega).

**Table S1.**
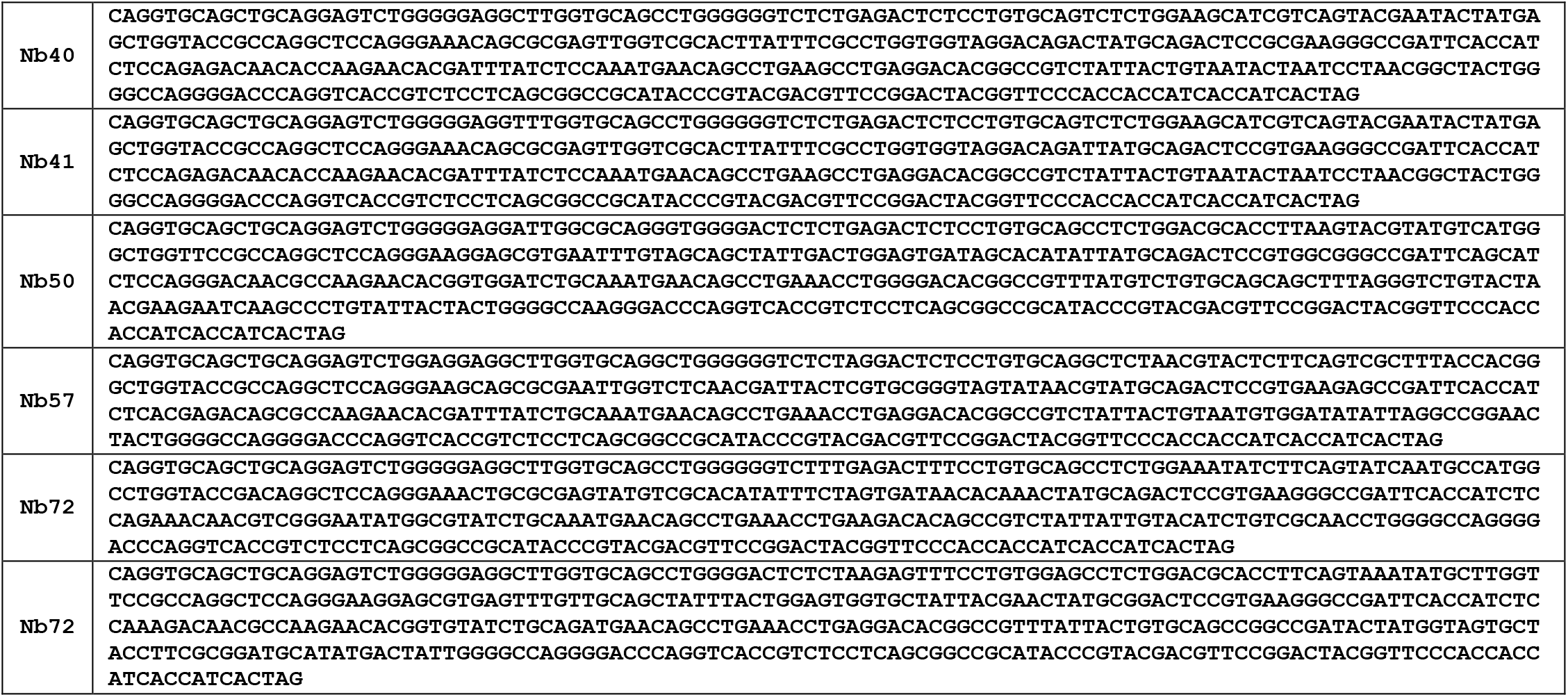
DNA sequences of the six laforin binding nanobodies.

**Figure S2.**
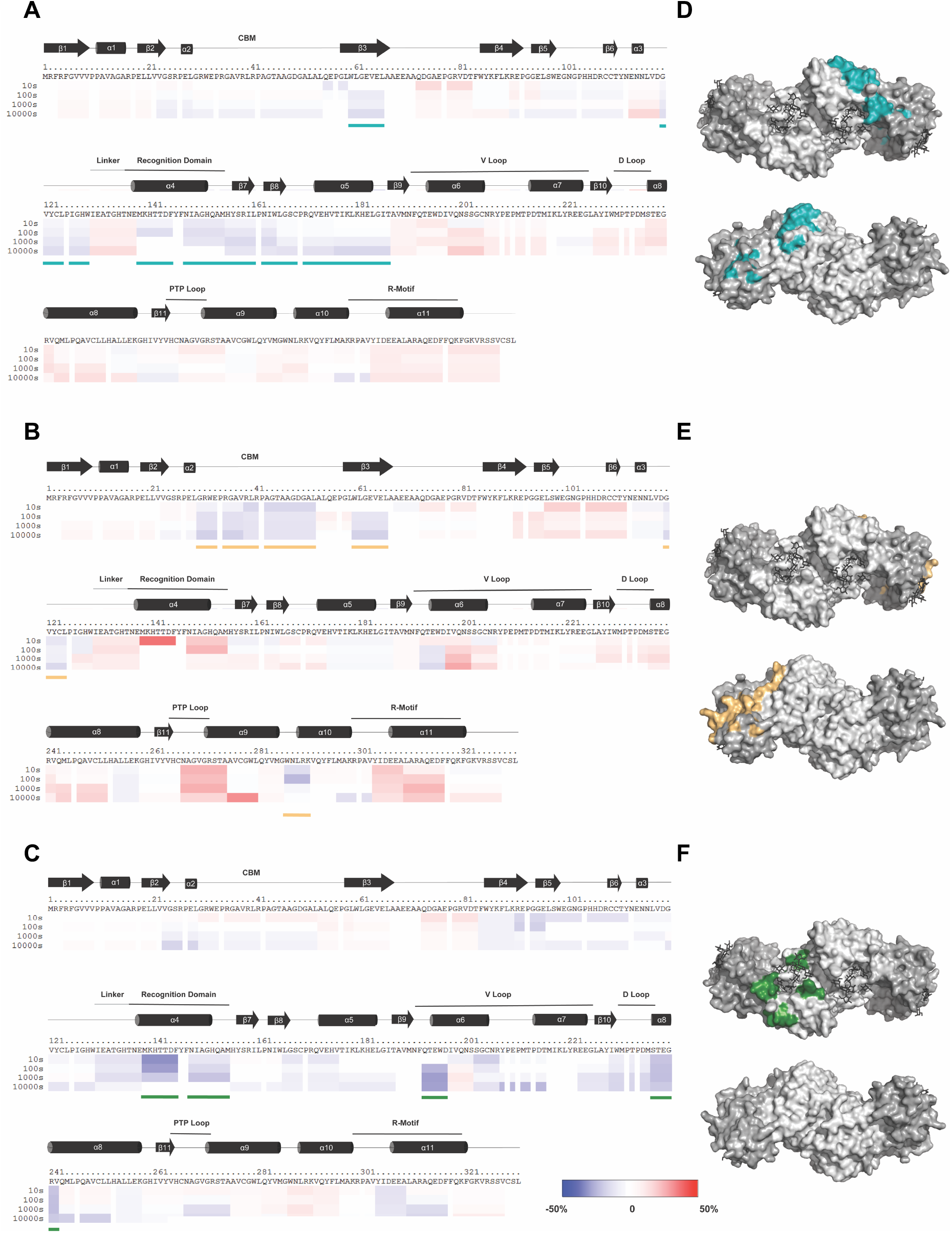
HDX analysis of anti-laforin nanobody binding. Fractions of each nanobody bound to laforin (Figure 3) were analyzed by hydrogen deuterium exchange mass-spectrometry (HDX) and the deuteration level was compared to laforin alone. The percent difference in deuteration between the nanobody complex versus laforin alone is represented with a negative percent change in blue to a positive percent change in red. Each colored bar below the primary sequence represents percent change in deuteration at one of four time points. Secondary structure elements are depicted above the primary sequence with the CBM and DSP motifs labeled. Regions where the nanobody caused a ≥15% change in deuteration are highlighted with a colored line. **A)** Laforin-Nb40 epitope spans β3, β7 and β8 pleated sheets and α4 and α5 helixes. Highlighted with a light teal line. **B)** Laforin-Nb50 epitope includes the β3 pleated sheet and partially covers the α10 helix. Highlighted with a light orange line. **C)** Laforin-Nb73 epitope covers the α4 helix, and partially covers the α6 and α8 helixes. Highlighted with a dark green line. **D,E,F)** The identified epitope for each nanobody was modeled onto the surface map of the laforin structure (PDB: 4RKK). Nb40 (light teal), Nb50 (light orange), and Nb73 (dark green) respectively mapped on laforin. The CBM and DSP of laforin are colored dark gray and light gray respectively.

**Figure S3.**
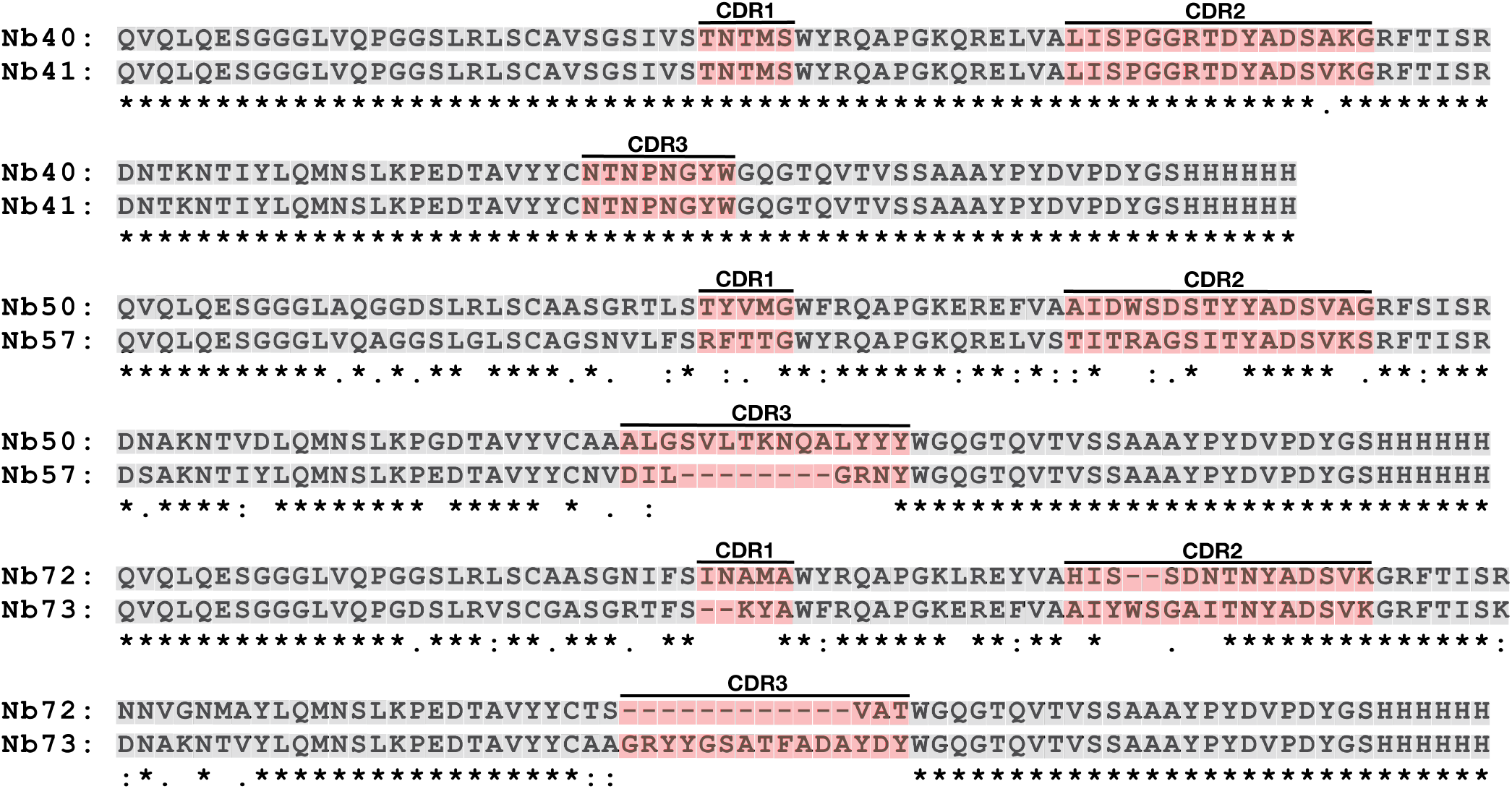
Anti-laforin nanobody sequence alignments. Sequence homology between Nb40 and Nb41 is 99.2%, Nb50 and Nb57 is 66.2%, and Nb72 and Nb73 is 70.6% (Clustal-Omega). CDRs highlighted in red, framework highlighted in gray.

**Figure S4.**
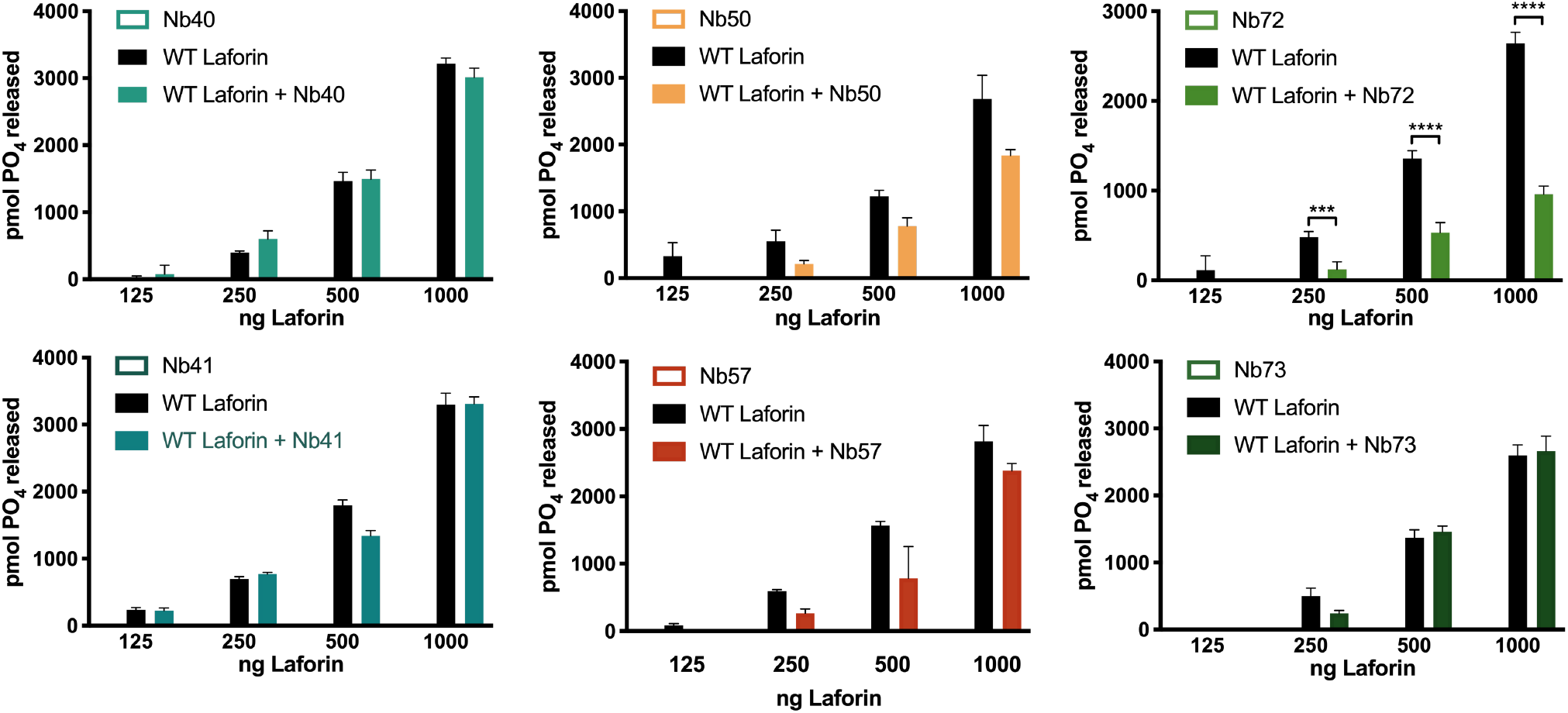
Laforin-nanobody activity assays. The specific activity of laforin against potato amylopectin in the presence and absence of nanobodies. Only Nb72 inhibits laforin’s phosphatase activity. Each bar is the mean ±SEM of 4 replicates. **** p < 0.0001.

